# Division of labor during biofilm matrix production

**DOI:** 10.1101/237230

**Authors:** Anna Dragoš, Heiko Kiesewalter, Marivic Martin, Chih-Yu Hsu, Raimo Hartmann, Tobias Wechsler, Carsten Eriksen, Susanne Brix, Knut Drescher, Nicola Stanley-Wall, Rolf Kümmerli, Ákos T. Kovács

## Abstract

Organisms as simple as bacteria can engage in complex collective actions, such as group motility and fruiting body formation. Some of these actions involve a division of labor, where phenotypically specialized clonal subpopulations, or genetically distinct lineages cooperate with each other by performing complementary tasks. Here, we combine experimental and computational approaches to investigate potential benefits arising from division of labor during biofilm matrix production. We show that both phenotypic and genetic strategies for a division of labor can promote collective biofilm formation in the soil bacterium *Bacillus subtilis*. In this species, biofilm matrix consists of two major components; EPS and TasA. We observed that clonal groups of *B. subtilis* phenotypically segregate into three subpopulations composed of matrix non-producers, EPS-producers, and generalists, which produce both EPS and TasA. This incomplete phenotypic specialization was outperformed by a genetic division of labor, where two mutants, engineered as specialists, complemented each other by exchanging EPS and TasA. The relative fitness of the two mutants displayed a negative frequency dependence both *in vitro* and on plant roots, with strain frequency reaching a stable equilibrium at 30% TasA-producers, corresponding exactly to the population composition where group productivity is maximized. Using individual-based modelling, we show that asymmetries in strain ratio can arise due to differences in the relative benefits that matrix compounds generate for the collective; and that genetic division of labor can be favored when it breaks metabolic constraints associated with the simultaneous production of two matrix components.

**Highlights:** - matrix components EPS and TasA are costly public goods in *B. subtilis* biofilms

- genetic division of labor using Δ*eps* and Δ*tasA* fosters maximal biofilm productivity

- Δ*eps* and Δ*tasA* cooperation is evolutionary stable in laboratory and ecological systems

- costly metabolic coupling of public goods favors genetic division of labor

## Introduction

Microbes can act collectively in groups, and thereby substantially influence their local environment to their own benefit. Such beneficial collective actions include the secretion of nutrient-degrading enzymes [1], iron-scavenging siderophores [2], biosurfactants for group motility [3], and structural components for biofilm formation [4,5]. In certain cases, cooperation even involves a division of labor, where subpopulations of cells specialize to perform different tasks [6,7]. Division of labor requires three basic conditions: individuals must exhibit different phenotypes (task allocation), the interaction between phenotypes must be cooperative, and all individuals must gain an inclusive fitness benefit from the interaction [8]. The allocation of tasks can be achieved either at the phenotypic or at the genotypic level. In the phenotypic specialization scenario, each individual carries genetic machineries for all tasks, but differences in gene expression result in tasks allocation [9]. In the genotypic specialization scenario, individuals carry only the genetic machinery for their own specialist task [9].

Both types of division of labor have been proposed to occur in microbes. For instance, during sliding colony expansion *Bacillus subtilis* cells phenotypically differentiate into surfactant producers and matrix producers where the role of the first is to reduce surface tension, while the latter allows expanding colony ‘arms’ to form and explore new territories [7]. Given the high relatedness between cells, specialization is likely beneficial for the group as a whole [8], with individuals gaining an inclusive fitness benefit from helping their clone mates [10–12]. However, division of labor has recently also been documented between genetically different strains or species [13,14].Cooperative division of labor based on genetic differentiation seems to evolve both frequently and reproducibly [14,15], lending support for the so-called Black Queen hypothesis, which depicts the microbial world as a network of interdependencies between species [16].

While our understanding of division of labor in microbes deepens [7,13,14], it has remained unclear what the advantages and disadvantages of the phenotypically vs genetically determined division of labor are, and which form yields higher fitness returns for the specialists and the community as a whole. When considering division of labor based on the exchange of two beneficial public goods, a phenotypic specialization could offer advantages because cells producing the two public goods will naturally be close to one another due to binary cell division. Close spatial proximity is essential for efficient public good sharing [18], yet might be compromised with genetically determined division of labor, as spatial separation of partners can readily occur and the switching of specialization states is not possible [19]. Conversely, genetically determined division of labor might offer advantages because it allows a complete decoupling of traits at the metabolic level. Specifically, the expression of two alternative synthetic pathways (both bearing a metabolic burden) can be terminally allocated into two different genetic lineages [20]. Finally, it has been argued that in contrast to phenotypic differentiation, terminal genetic divergence bears risks of conflicts such as social exploitation because relatedness between interacting partners is reduced; potentially leading to diverging interests between partners [19].

Here, we focus on identifying the costs and benefits associated with different division of labor strategies for biofilm formation in the common soil and plant-colonizing bacterium *B. subtilis*, in terms of individual and collective productivity. Biofilms represent the most common lifestyle of bacteria, where cells are in close proximity to one another, embedded in extracellular matrix (ECM) [21]. There is ample opportunity for division of labor over matrix construction, because ECM usually consists of multiple secreted compounds that form a mesh of complex exopolysaccharides (EPS) and structural proteins, sometimes accompanied by extracellular DNA (eDNA). While the presence of eDNA can be the consequence of cell death [22], the production of matrix exopolysaccharides and proteins tends to be triggered by cooperative signaling [23], cues released by competitors [24], or specific nutrient components [25,26]. As the synthesis of large polymers is metabolically costly, tight regulation of matrix gene expression is often in place, and it has been suggested that the overall metabolic costs for the community may be reduced by assigning matrix production only to a subpopulation of cells [27]. Here we propose an alternative scenario involving division of labor, where subgroups of individuals within a biofilm each specialize (either phenotypically or genetically) in the production of a different matrix component, which are then shared at the level of the group.

Our model system involves *B. subtilis* forming robust, wrinkly pellicle biofilms that reside at the oxygen rich liquid-air interface [28]. Increasing cell density of the planktonic cells results in a decreasing oxygen concentration in the bottom layers of the static medium. Aerotaxis of *B. subtilis* leads to an accumulation of cells near the liquid-air interface and eventually a colonization of the surface in a form of a densely packed pellicle biofilm. During pellicle development transcription of the matrix-related operons *epsA-O* and *tapA-sipW-tasA* is derepressed [27,29–31] eventually allowing synthesis of the biofilm exopolysaccharide (EPS), and the structural protein TasA [32,33]. Mutants lacking either EPS or TasA cannot establish pellicle biofilms individually, but they can complement each other in co-culture, indicating that both matrix components are necessary for pellicle biofilms and that they are shared [32,34,35].

Using a mixture of fitness assays, single-cell gene expression analyses and mathematical modelling, we show that the two matrix components EPS and TasA are indeed costly to produce. We further found that cells within a biofilm phenotypically differentiate into three distinct subpopulations consisting of cells producing either both of the matrix components, EPS alone, or none of the two components. We then demonstrate that in terms of group productivity, genetic division of labor for matrix construction can be superior to the phenotypic differentiation strategy present in the wild type. Specifically, biofilm productivity was maximized at an intermediate mixing ratio of mutants deficient for either EPS or TasA, both in pellicle biofilms grown in the laboratory, and in biofilms grown on plant hosts. Crucially, the Δ*eps*: Δ*tasA* proportion at which biofilm productivity maximization occurred, represents a stable equilibrium.

## RESULTS

### The matrix components EPS and TasA serve as costly public goods

Components of bacterial extracellular matrix are often large, complex polymers, which can potentially bear significant metabolic production costs [1,36]. To demonstrate the costs associated with the production of EPS and TasA in our *B. subtilis* strain (NCBI 3610), we competed the non-producing mutants Δ*eps* and Δ*tasA* against the wild type (WT) under conditions where matrix is synthesized but not required for survival [37], which is up to 16 hours of growth, prior to surface colonization (Movie S1; see Methods). We confirmed that in the pre-pellicle phase the WT, Δ*eps*, and Δ*tasA* strains first grow exponentially before reaching the early stationary phase (Figure S1A). While strains expressed the corresponding matrix components (Figure S1B-D assays based on fluorescent transcriptional reporters P_*eps*_*-gfp* and P_*tapA*_*-gfp*), the expression patterns slightly varied between the WT and the mutants. The expression of P_*tapA*_ in Δ*eps* and P_*eps*_ in Δ*tasA* were slightly stronger and weaker, respectively, with shift towards more homogenous expression in both mutants (Figure S1C). Under these conditions, our growth competition fitness assay revealed significant costs for both matrix components (Figure 1A). The fact that Δ*eps* had significantly higher relative fitness than Δ*tasA* in pairwise competition against the WT suggests that EPS synthesis bears a higher cost than TasA production under these conditions (Figure 1A).

**Figure 1.**
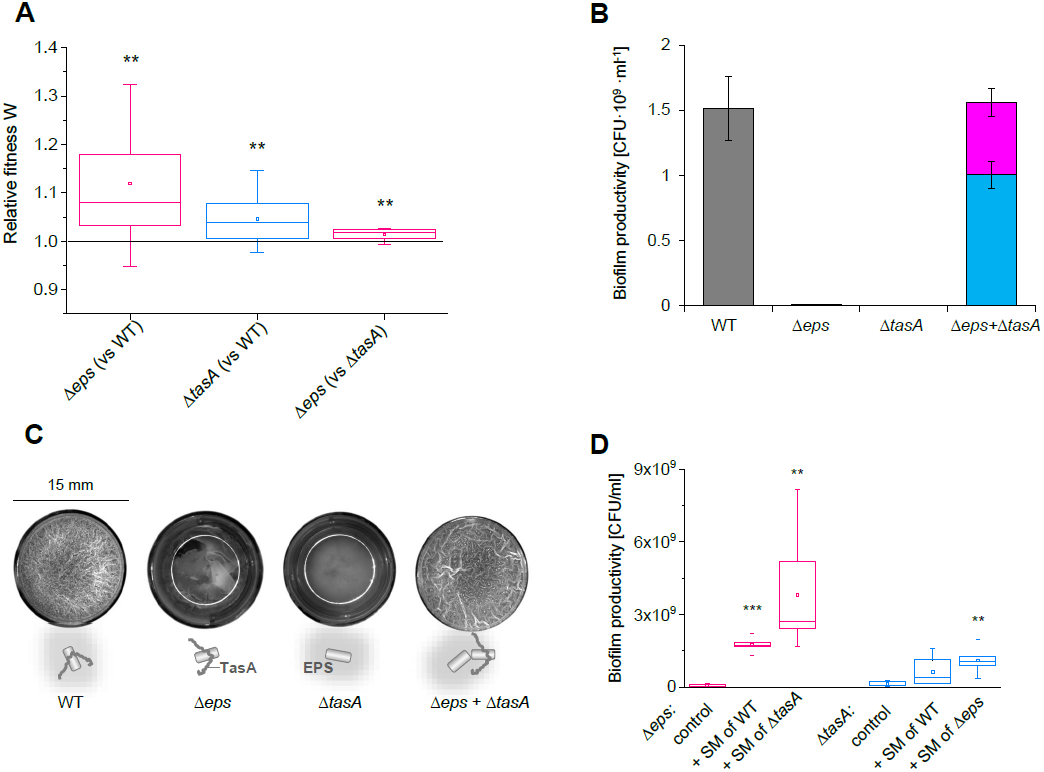
Costs and benefits of matrix components EPS and TasA. **(A)** To estimate the metabolic costs of EPS and TasA production the matrix-deficient strains Δ*eps* and Δ*tasA* were competed against the WT and against each other under conditions where matrix components are produced but not required (see Methods). Relative fitness (W) was calculated for Δ *eps* (when competed against WT), Δ*tasA* (when competed against WT), and Δ *eps* (when competed against Δ*tasA*). Relative fitness W significantly larger than 1 indicates an advantage of a given strain in a given pair-wise competition. For Δ *eps* vs WT n=13, p<003; for Δ*tasA* vs WT n=13, p<008; for Δ*tasA* vs WT n=8, p<007 (**p<0.01, *** p<0.001). **(B)** Productivity of the WT, Δ*eps*, Δ*tasA* and Δ *eps* +Δ*tasA* co-culture (50:50 ratio) measured as CFU/ml. Data points represent mean and error bars represent standard error obtained from biological triplicates. **(C)** Brightfield images of pellicle morphology developed by the WT, matrix-deficient mutants in monocultures and by the Δ*eps*+Δ*tasA* co-culture (50:50 ratio). The cartoons below represent public goods produced by each culture. **(D)** To confirm that EPS and TasA can be shared and thereby serve as public goods, the matrix-deficient strains Δ*eps* and Δ*tasA* were allowed to form pellicles in presence of spent media (SM) obtained from the WT or from the complementary mutant (n=4-6). Pellicle productivity [CFU/ml] reached in presence of those SMs was compared with the productivity of the control (a strain exposed to its own SM): for Δ*eps* + SM of WT p<5×10^−7^; Δ*eps* + SM of Δ*tasA* p<0.008; Δ*tasA* + SM of WT p<; Δ*tasA* + SM of Δ*eps* SM p<0.001 (**p<0.01). Boxes represent Q1–Q3 (quartiles), lines represent the median, and bars span from max to min. To better distinguish between the matrix-deficient mutants, data for Δ*eps* and Δ*tasA* are presented in pink and blue, respectively.

Next, we examined sharing of the two components. We began with complementation assays mixing the two mutants (Δ*eps* and Δ*tasA* single deletion mutants) in 1:1 ratios. In line with previous reports [32,34,35], we found that the mutants could not establish pellicles when grown in monocultures, but complemented each other when co-cultured, indicating that EPS and TasA can be shared (Figure 1B,C). Since TasA was previously depicted as a cell-associated amyloid fiber, anchored through the accessory protein TapA to the cell [34], we performed additional experiments to confirm cross-complementation. Specifically, we added conditioned media from the EPS and TasA producers to growing cultures of the Δ*eps* and Δ*tasA*, respectively, and quantified their surface colonization ability. We observed that the conditioned medium from the WT or the complementary mutant significantly improved pellicle formation as compared to the control, with the effect being more pronounced for the Δ*eps* than the Δ*tasA* mutant (Figure 1D). This result suggests that the spent medium obtained from the WT and Δ*tasA* contained freely released EPS which could complement the Δ*eps* strain. Similarly, WT and Δ*eps* released a portion of TasA into the medium, that could partially complement the Δ*tasA* phenotype.

As the above results (Figure 1D) suggest that the matrix components EPS and TasA differ in the extent to which they are shared, pointing towards stronger privatization of TasA, we hypothesized that efficient mixing of EPS-producers and TasA-producers is necessary for successful complementation. To test the role of mixing we took advantage of a previously observed motility effect on cell assortment in pellicle biofilms [38]: Cells lacking a functional flagellum (Δ*hag*) are less efficient in swimming to the top of the liquid, which likely results in very low number of founder cells carrying the Δ*hag* mutation (compared to WT) in the pellicle. As a result, pellicles formed by two isogenic Δ*hag* strains labeled with different fluorophores, contain large clusters of cells of the same lineage, indicating limited genotype mixing [38]. As expected, the efficiency of complementation between EPS- and TasA-producers was negatively affected in the Δ*hag* genetic background as compared to the control with functional flagella (Figure S2A). Finally, the spatial assortment of cells in the pellicles formed by mixtures of Δ*eps* and Δ*tasA* and pellicles formed by the WT were compared using a density correlation function quantification method (see Methods), to assess the spatial effects of genetic division of labor (Figure S2B-D). The level of spatial strain mixing was slightly higher in pellicles formed by mixtures of Δ*eps* and Δ*tasA* (regardless of the fluorescence reporter combination) as compared to pellicles formed by the WT (Figure S2C,D).

Altogether, these results confirm that both matrix components EPS and TasA can be shared and that robust pellicle biofilm formation depends on the efficient exchange of these compounds.

### Wild type cells exhibit phenotypic heterogeneity in the expression of matrix components

As EPS and TasA are costly to produce (Figure 1A) and can both be shared between the producers and non-producers (Figure 1B-D), we hypothesized that phenotypic differentiation into EPS-producers and TasA-producers could occur and form the basis of a division of labor in WT *B. subtilis* populations during pellicle formation. To test for phenotypic heterogeneity of *eps* and *tasA* expression at the single cell level, we used a reporter strain carrying a promoter fusion of the *eps* promoter to *gfp* (P_*eps*_*-gfp*) and an analogous reporter for the *tapA* promoter based on *mKate* (P_*tapA*_*-mKate*) at two distinct genomic loci (see Methods, Table S1). As a control, we used the P_*tapA*_*-gfp* P_*tapA*_*-mKate* strain (see Methods, Table S1) for which no phenotypic heterogeneity and a linear correlation between the two fluorescence channels was expected. Fluorescent images of mature pellicles of the WT P_*eps*_*-gfp* P_*tapA*_*-mKate* strain and the control WT P_*tapA*_*-gfp* P_*tapA*_*-mKate* strain were captured using confocal laser scanning microscopy (CLSM). While the control strain showed a clear spatial correlation between GFP and mKate fluorescence intensities, this was not the case for the WT P_*eps*_*-gfp* P*tapA-mKate* strain (Figure 2A). Specifically, large bright clusters of strong GFP signal could be observed in locations in which there was reduced mKate fluorescence, suggesting the presence of a subpopulation that is partially specialized for EPS production (Figure 2A). We further performed quantitative analyses of the fluorescent images by artificially dissecting the images into small cubes and quantifying the GFP- and mKate-signal intensities in each cube (see STAR Methods). This allowed us to examine whether GFP and mKate fluorescence intensities linearly correlate in space. Such linear correlation was expected from the control strain (WT P_*tapA*_*-gfp* P_*tab*_*-mKate*) and could be the case for the P*eps-gfp* P_*tapA*_*-mKate* strain if *eps* and *tasA* were expressed by the same population of cells. The analysis confirmed that for biofilms made by the P_*tapA*_*-gfp* P_*tapA*_*-mKate* strain, signal intensities from GFP and mKate channels showed strong linear correlation in space, this correlation was much weaker in case of the P_*eps*_*-gfp* P_*tapA*_-*mKate* strain (Figure 2B, Figure S3A).

**Figure 2.**
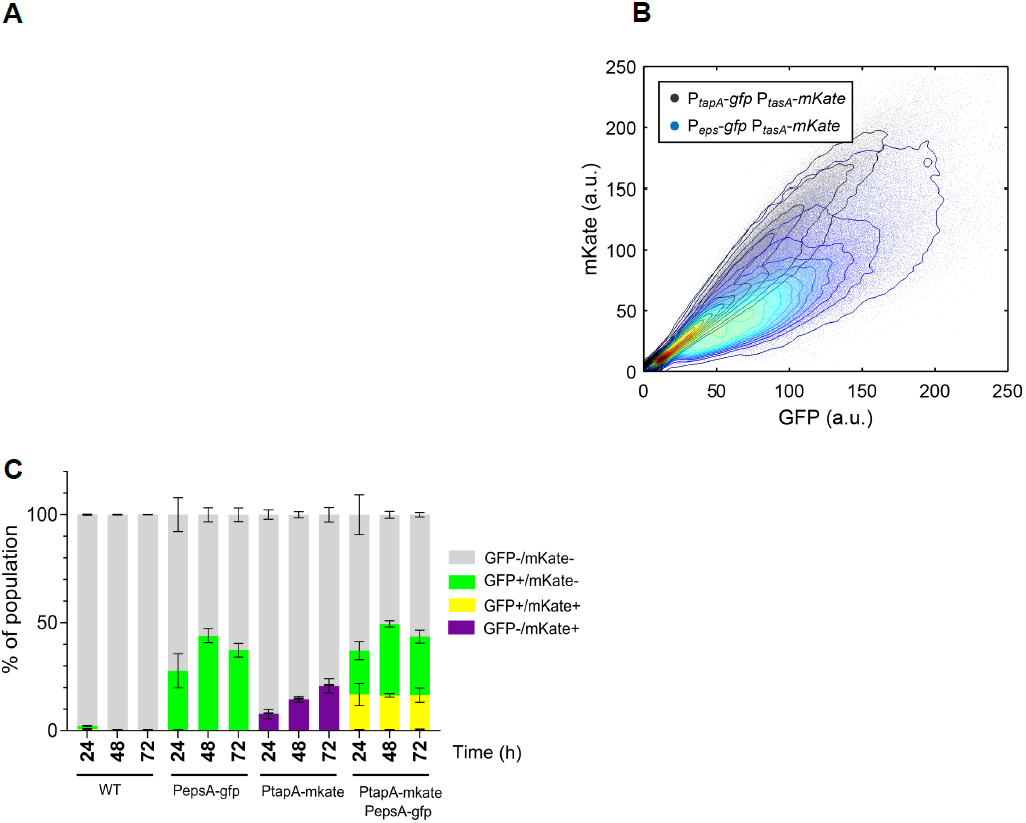
Native phenotypic heterogeneity in the expression of matrix components. **(A)** Pellicles formed by the double-labelled strain carrying the P_*eps*_*-gfp* P_*tapA*_*-mKate* reporters and the strain carrying the P_*tapA*_*-gfp* P_*tapA*_*-mKate* reporters (control) were visualized using a confocal microscope to compare the distribution of fluorescence signal from different fluorescence reporters (GFP, mKate). **(B)** Volumes in GFP and mKate fluorescence channels (obtained by manual thresholding) were merged, dissected into cubes and the average intensities in the GFP and mKate channels for all cubes were plotted (see Methods). The maximum density is normalized to 1 and the contour lines correspond to 0.05 decrease in density. **(C)** The following strains: NCIB3610, NRS2242 (carrying P_*eps*_*-gfp*), NRS3913 (carrying P_*tapA*_*-mKate*), and NRS5832 (carrying P_*eps*_*-gfp* and P_*tapA*_*-mKate*); were allowed to form pellicles and were then analyzed using flow cytometry. Bar chart (mean±SD) represents fraction of OFF cells, cells expressing *eps-gfp, tapA-mKate*, and cells expressing both reporters (n=3).

The above experiment suggests that matrix-expressing subpopulations of WT *B. subtilis* exhibit a certain degree of phenotypic differentiation into cells that produce mostly EPS and cells that produce both EPS and TasA (generalists). To confirm this pattern, we analyzed single cells extracted from pellicles using fluorescence-guided flow cytometry (FC). FC analyses were performed at 3 time points during pellicle development (24, 48 and 72 hours) and included controls with strains carrying single reporter fusions (see Methods, Table S1). These analyses revealed the presence of 3 distinct subpopulations of cells: (i) matrix-OFF cells where fluorescence signals from both the P_*eps*_ and P_*tapA*_ promoters were below the detection thresholds; (ii) matrix-ON cells where there was a positive linear correlation of the signals from the P*eps* and P_*tapA*_ promoters: (iii) EPS-ON cells, containing a fluorescent signal from P_*eps*_, but not from P_*tapA*_ (Figure 2C, Figure S3B). Differences in relative frequencies of P_*eps*_*-gfp* and P_*tapA*_*-mKate* ON cells were not due to the use of different fluorescent reporters, as evidenced by our FC control experiments where strains carrying either a P_*tapA*_*-gfp* or a P_*tapA*_*-mKate* reporter construct showed identical frequencies of ON cells (Figure S3C,D). Thus, our FC experiments confirmed that the expression of the two major matrix promoters P_*eps*_ and P_*tapA*_ is not perfectly correlated, which likely translates into phenotypic diversity at the level of EPS and TasA production in wild type *B. subtilis* pellicles.

### Genetic division of labor yields higher biofilm productivity than phenotypic differentiation

Although the above data indicate that wild type cells differentiate into EPS-producers, generalists, and non-producers during pellicle biofilm formation, this pattern does not resemble the canonical principle of division of labor where distinct subpopulations of cells are expected to either commit completely to TasA or EPS production. The incomplete specialization could be due to regulatory constraints. For instance, it is known that the *epsA-O* and *tapA-sipW-tasA* operons share multiple regulators, suggesting that some level of parallel expression (either on or off) at the single cell level is expected [39,40].

We thus wondered whether an incomplete specialization represents a beneficial strategy or whether it can be outperformed by a genetically determined specialization, where cells are ultimately constrained in the production of either TasA or EPS. To address this question, we studied the division of labor between Δ*tasA* as the exclusive EPS-producer and Δ*eps* as the exclusive TasA producer. In a first experiment, we mixed the exclusive EPS-and TasA-producers at different ratios and examined the productivities of pellicles (Figure 3A). We found that pellicle productivity varied in response to strain frequency, and peaked at a strain ratio of approximately 30 % Δ*eps*: 70% Δ*tasA* (Figure 3A). Interestingly, the group productivity of mixtures close to this optimal ratio was significantly higher than the WT productivity, indicating that the genetic division of labor over matrix construction outperforms the native phenotypic differentiation observed in the WT (Figure 3A).

**Figure 3.**
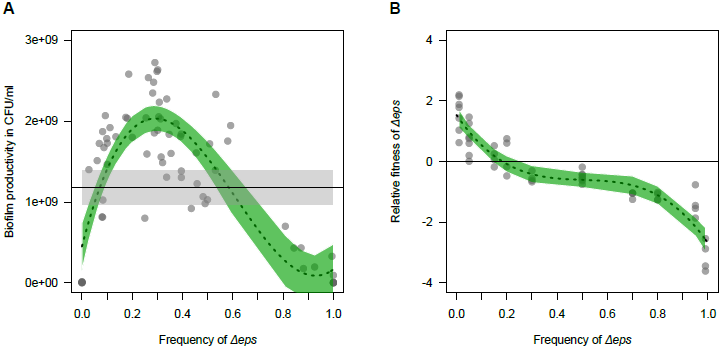
Productivity and fitness in pellicles with genetic division of labor. **(A)** Productivities of Δ*eps*+Δ*tasA* biofilms [CFU/ml] measured for different mixing ratios and compared to average productivity reached by the WT (black horizontal line with grey shaded 95 % confidence interval). The dashed line and green shaded 95% CI represent a cubic fit to the fitness data (F_3, 68_ = 54.9, R^2^ = 0.695, p < 0.0001). **(B)** The relative fitness of Δ*eps* in competition with Δ*tasA* followed a negative-frequency dependent trajectory, best described by a cubic fit (dashed line with 95 % CI: F_3,46_ = 94.7, R^2^ = 0.852, p < 0.0001).

### Genetic division of labor is evolutionarily stable in pellicles and on plant roots

We next asked whether such genetic division of labor, which yields the highest fitness returns at a strain ratio of approximately 30:70, represents a stable equilibrium or simply a transient phenomenon. To test this possibility, we competed the Δ*eps* strain against the Δ*tasA* strain across a range of frequencies (1% to 99 %), over the full cycle of pellicle growth (from inoculation until formation of robust, wrinkly pellicle after 48 hours). These competitions revealed that the relative fitness of Δ*eps* followed a negative frequency-dependent pattern: Δ*eps* outcompeted Δ*tasA* when rare, but lost the competition when common (Fig. 3B). Strikingly, the two strains showed equal competitiveness at starting frequencies between 20% - 30% Δ*eps*, thus exactly at the strain ratio where biofilm productivity is maximized. These findings strongly suggest that, regardless of the metabolic cost imbalance between the two matrix components, stable coexistence of the EPS and TasA producers is favored in the pellicle, with strain frequency evolving towards the optimum in terms of biofilm biomass productivity (Figure 1A, Figure 3B).

To test whether stable genetic division of labor could also manifest in a relevant ecological environment, we repeated several key experiments using plant root associated biofilms. Specifically, we subjected the roots of *Arabidopsis thaliana* seedlings to colonization by the WT, or a mixture of Δ*eps* and Δ*tasA* strains at a 50:50 ratio, or monocultures of the two mutants (see Methods). Each strain carried a constitutive fluorescent reporter to allow biofilm visualization by CLSM (see methods, Table S1). In line with previous studies [41], both the WT and the mixture of Δ*eps* and Δ*tasA* strains were able to produce thick biofilms on the roots, which was not the case for the Δ*eps* and Δ*tasA* mutants grown in monocultures on the plant root (Figure 4A,B). Analogous to the pellicles, we found that the productivity of root biofilms was significantly higher for the Δ*eps* + Δ*tasA* mixture as compared to the WT. Next, we estimated the relative frequencies of Δ*eps* and Δ*tasA* mutants in the mixed biofilm on the root, based on total pixel volumes (see methods), and found that the mutant frequency settled at the optimal ratio of 20% - 30% Δ*eps* (Figure 4B,C). In contrast, the frequency remained close to 0.5:0.5 in our control mixtures of two WT strains labeled with different fluorescent reporters (Figure 4B,C). Altogether, our experiments demonstrate that the genetically hard-wired division of labor between EPS- and TasA-producers provides fitness benefits not only in pellicles, but also on plant roots.

**Figure 4.**
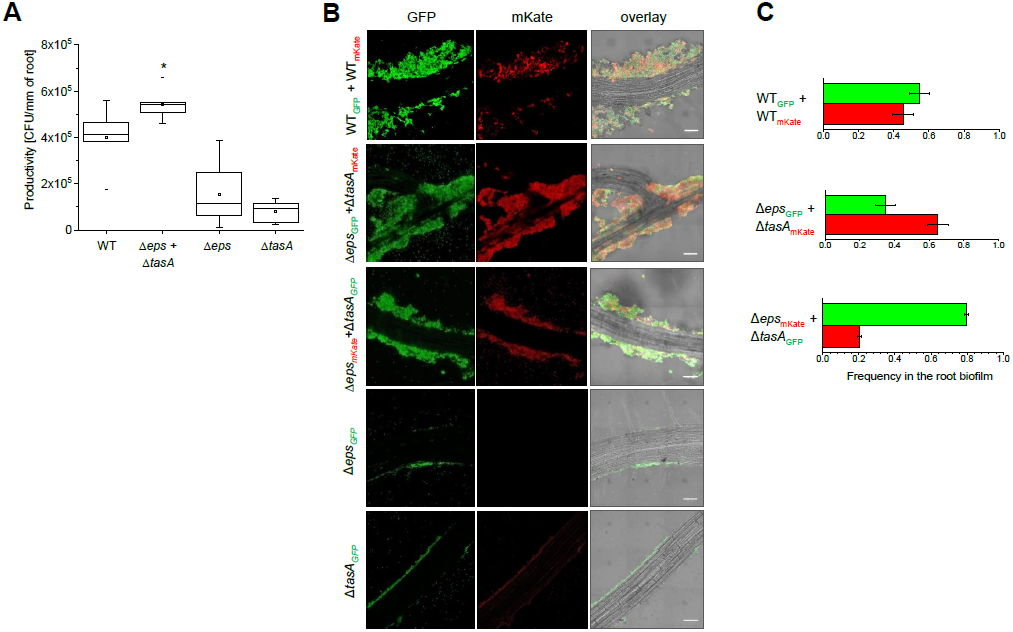
Genetic division of labor on plant roots. **(A)** *Arabidopsis thaliana* roots were colonized by the WT, Δ*eps*+Δ*tasA* mix and the mutants in monocultures and biofilm productivities were measured as CFU/mm of root (for WT and co-culture n=6; for Δ*eps* and Δ*tasA* n=11) (see Methods). The productivity reached by the Δ*eps*+Δ*tasA* mixture was compared with productivity of the WT (p<0.04).**(B)** *A. thaliana* roots were colonized by mixed cultures of WT_GFP_+WT_mKate_,Δ*eps*_GFP_+Δ*tasA*_mKate,_Δ*eps*_mKate_+Δ*tasA*_GFP_ and the mutants in monocultures (Δ*eps*_GFP_ and Δ*tasA*_GFP_), and visualized using CLSM. Scale bar represents 10μm. **(C)** Frequencies of each strain in the root-associated biofilm were determined based on image analysis (see Methods). Bars represent average (n=3-5) and error bars represent standard error.

### Simulating pellicle formation to understand the drivers of genetic division of labor

To better understand the conditions required for genetic division of labor to evolve between EPS- and TasA-producing specialists, we used an individual-based modelling platform, specifically developed to simulate microbial interactions [42]. The platform consists of a two-dimensional toroidal surface, where bacterial cells are modeled as discs. Bacteria are seeded in low numbers to their *in-silico* habitat, and are then allowed to consume resources, grow, divide, disperse, and produce public goods according to specified parameters for 10,000 time steps (see methods for fitness equations). The system keeps track of both bacterial strains and their public goods over time and space, and closely recovers patterns of real pellicle formation (Figures S4).

First, we examined the performance of a wildtype (WT) strain, which simultaneously produces two complementary public goods, representing EPS and TasA. Simulations started with eight cells placed in the center of the landscape to mimic the early phase of pellicle formation. Cells grew and divided at a basic rate (μ). Cells additionally produced diffusible public goods at a constant rate. Public goods diffuse randomly, can decay or generate fitness benefits for receiver cells. While each public good generates a benefit on its own, synergistic benefits accrue to cells that encounter the two complementary public goods within a certain time frame. Using this setup, we found that WT biofilm productivity peaked with lower public good diffusion *d* (Figure S5A), indicating that reduced diffusion minimizes the loss and improves sharing of public goods. Since our experimental data suggest that TasA and EPS differ in the level of sharing, and thus in the relative benefit these goods can generate for the group, we varied this parameter in our model, but found that it did not affect the productivity of WT pellicles (Figure S5B). We then implemented metabolic constraints (via the factor *f*, defined in Eq. (1) in the STAR Methods) in the WT to account for the possibility that the simultaneous production of two public goods exceeds the sum of each individual public good (*f* > 1) [20]. We observed that biofilm productivity sharply declined with increased levels of constraints (Figure S5C).

Next, we considered different levels of phenotypic heterogeneity in the WT by starting simulations with different ratios of specialist and non-specialist cells (Figure S5D-F). We found that any level of phenotypic heterogeneity outperformed uniform trait expression at the beginning of pellicle formation (Figure S5D). Conversely, the most beneficial strategies in more mature pellicles were either complete specialization or no specialization at all (Figure S5E,F). We hypothesize that no specialization performs well because it allows efficient public good sharing at higher cell densities, and complete specialization is beneficial because it breaks metabolic constraints. In contrast, any intermediate form of phenotypic heterogeneity suffers from reduced sharing and sustained metabolic constraints, and should thus be selected against. This finding might explain why the *B. subtilis* WT strain showed relatively low levels of phenotypic heterogeneity.

We then asked whether two genetically fixed mutants, producing either one or the other public good (i.e. mimicking Δ*eps* and Δ*tasA* mutants) can complement each other. Similar to our empirical observations, we found successful complementation between the two specialist strains, with pellicle productivity peaking at intermediate mixing ratios (Figure 5A). Moreover, the relative fitness of the Δ*eps* strain exhibited negative-frequency dependence (Figure 5B), and the point of fitness equilibrium occurred exactly at the productivity peak of the group. Next, we implemented the experimental observation that TasA yields lower benefits than EPS. We again observed successful complementation, but pellicle productivity reached higher levels and peaks shifted to lower frequencies of the Δ*eps* strain, in the case of a greater benefit imbalance between the two public goods (Figure 5C). The relative fitness of the Δ*eps* strain again followed a negative-frequency dependence with the point of intersection being exactly at the pellicle productivity peak (Figure 5D). To examine whether the reciprocal symmetric exchange of public goods is the reason for why strain equilibrium frequency coincides with maximal group productivity, we simulated a scenario of asymmetrical public good exchange between a strain producing both public goods and a strain producing a single public good (Figure 5E,F). For this scenario, we found that the relationship between strain equilibrium and maximal group productivity breaks: the strain producing only one public good experienced relative fitness advantages at all strain frequencies, driving strain frequency away from maximal group fitness.

**Figure 5.**
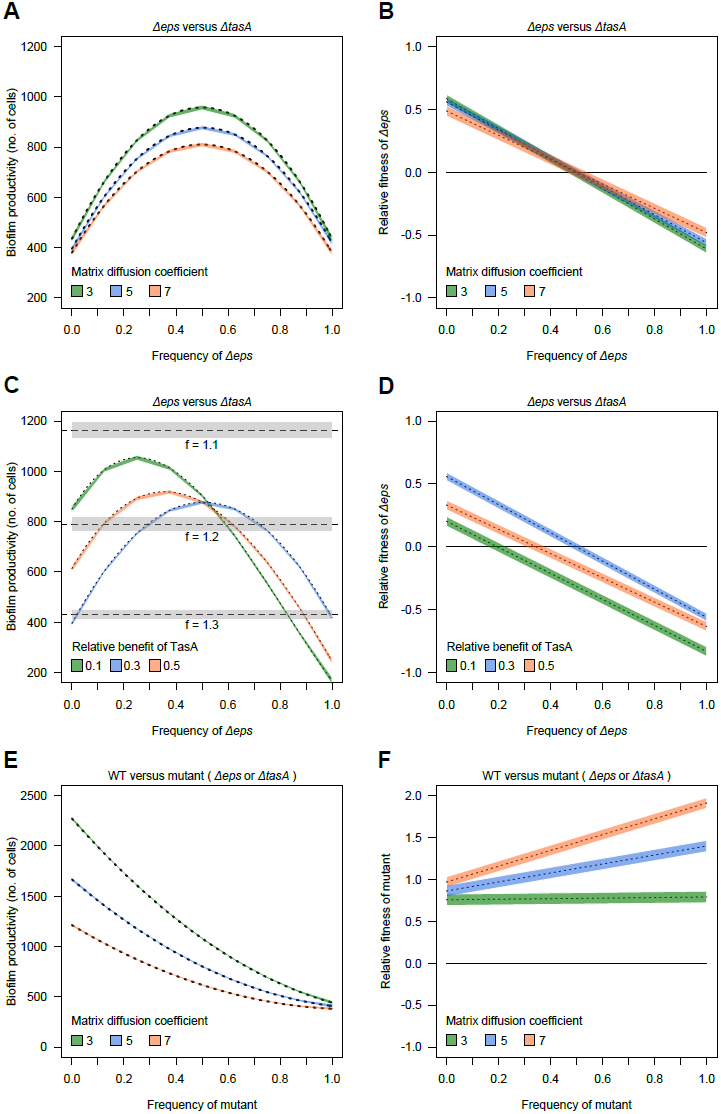
Individual-based simulations identify drivers of genetic division of labor. We simulated biofilm formation of the mutants Δ*eps* (producing TasA) and Δ*tasA* (producing EPS) when grown in co-culture. Biofilms were initiated with eight cells, with Δ*eps* frequency varying between 0 and 1, in steps of 0.125. Cells produced diffusible matrix components (either TasA or EPS) and grew according to their fitness functions. After 10,000 time steps, we measured the absolute productivity of the biofilm (no. of cells) and the relative fitness of the competing strains within biofilms. Fitness trajectories are shown as the best fit from linear models across 50 simulations for each condition (± 95 % confidence interval).**(A)** and **(B)** depict variation in biofilm productivity and relative fitness of mutants, respectively, as a function of strain frequency and the matrix diffusion coefficient, under conditions where both matrix components generate equal benefits. **(C)** and **(D)** show variation in biofilm productivity and relative fitness of mutants, respectively, as a function of strain frequency and different relative benefits of the two matrix components (diffusion coefficient *d* = 5). Dashed lines and grey shaded area in (**D)** depict mean ± 95 % CI productivities of WT biofilms across a range of metabolic constraints (*f*) accruing in the WT for simultaneously producing two public goods. **(E)** and **(F)** depict biofilm productivity and relative fitness of a mutant growing together with the WT, respectively, as a function of strain frequency and the matrix diffusion coefficient, under conditions where both matrix components generate equal benefits.

Finally, we asked whether genetic division of labor between the Δ*eps* and Δ*tasA* strains can outperform the WT strategy, as observed in our empirical experiments. However, in the absence of any metabolic constraints (*f* = 1), the WT pellicle productivity was 1714 ± 39 cells (mean ± SE, with intermediate diffusion *d* = 5), and thus by far higher than productivity in any of the complementation scenarios (Figure 5). Conversely, when the WT faces metabolic constraints, we found a parameter space (*f* > 1.1), in which pellicle productivity of complementing strains exceeds wildtype performance (Figure 5C).

Taken together, our simulations recover the key features of our experimental system and suggest that the reciprocal symmetrical exchange of public goods under conditions of relatively low diffusion together with the decoupling of metabolic constraints are the preconditions required for the evolution of stable genetic division of labor over biofilm matrix production.

## DISCUSSION

Despite their unicellular simplicity, microbes can coordinate complex behaviors as a group. Some of these multicellular behaviors involve division of labor between phenotypically distinct subpopulations [7,17], or even different genetic lineages [14]. Here we deployed a combination of experiments and simulations to directly compare these two alternative cooperative strategies. By focusing on the production of two biofilm matrix components in *B. subtilis*, we found evidence for significant, yet incomplete phenotypic specialization in matrix production among clonal cells of the wild type strain. However, this strategy of phenotypic specialization was outperformed by genetic division of labor, where strains, engineered as strict specialists, settled on an equilibrium ratio that maximized biofilm productivity. Our individual-based modeling approach captures the experimental system and reveals that metabolic decoupling of two costly traits can be the key to success for genetic specialization.

While we demonstrate that *B. subtilis* WT displays partial phenotypic differentiation at the level of matrix production, we might ask why this form of specialization is not more pronounced, especially in the context of the reported fitness benefits that can accrue from complete genetic specialization (Fig. 3A). One explanation might be that the *epsA-O* and *tapA-sipW-tasA* operons share multiple regulators, such that some level of parallel expression is inevitable [39,40,43,44]. Still, there could be certain mechanisms in place to decouple EPS and TasA production, for example a positive feedback where EPS prevents autophosphorylation of the EpsAB kinase, allowing activation of the EpsE glycosyl-transferase, thereby promoting EPS-synthesis [45]. It was also proposed that the major matrix repressor SinR, acts differently on P_*eps*_ and P_*tapA*_ promoters. Specifically, in the case of P_*eps*_ it directly competes with an activator RemA for the binding site upstream of the promoter, thereby serving as an anti-activator, while in case of P_*tapA*_ it binds simultaneously with RemA, probably serving as a repressor [44]. The opposing relationship between SinR and RemA may lead to an outburst of *epsA-O* expression in a subpopulation of cells, while *tapA-sipW-tasA* remains under tighter control of SinR. While these regulatory mechanisms could allow for some heterogeneity in gene expression, complete specialization seems impossible. Another possibility is that such complete specialization with 30:70 frequencies of TasA-producers and EPS-producers, does not always reflect an optimal solution. Biofilm formation is only one out of multiple cooperative survival strategies of *B. subtilis* and each strategy might require different optimal ratios of generalists and specialists. For instance, the presence of generalists that can produce both EPS and TasA may be favored during social spreading on solid surfaces, where both components are important [7], but where the diffusion of these public goods is reduced compared with pellicle growth conditions.

Our findings on successful genetic division of labor between specialized strains, producing either EPS or TasA, show that strain frequency settles as a stable frequency of approximately 70:30. Our model suggests that the dominance of EPS-producers in biofilms may be driven by a higher relative benefit of EPS compared with TasA. In addition, the altered matrix gene expression patterns in Δ*eps* (overproducing TasA) and Δ*tasA* (reduced EPS production) suggest that a higher proportion of Δ*tasA* might be required for stable pellicle production. Perhaps the two matrix components engage in an interaction that alters their biochemical properties. One possibility is that upon interaction with EPS, TasA becomes less soluble and vice versa. This could explain a decreased complementation efficiency of Δ*eps* and Δ*tasA* strains by the spent media obtained from the WT (Figure 1D). Recent work suggests that the structural functionality of TasA fibers may directly depend on the presence of EPS in the extracellular environment [46].

The 70:30 population structure is stable in typical laboratory setup (pellicle biofilms) and in plant root-associated biofilms. Although the genetic division of labor arose as the winning strategy, our study also points towards the canonical problem associated with fixed cooperation strategies: limited mixing of strains prevents efficient genetic division of labor [19]. Specifically, we found that the complementation between EPS- and TasA-producers was ineffective in experiments with flagellum-deficient strains, which exhibit a decreased level of mixing, thereby reducing public good sharing and the formation of robust pellicle biofilms. Mathematical models ([47], our model) suggest that complementation is most efficient when strain mixing is high, but the diffusion of public goods is reduced, conditions that foster efficient public good exchange between neighbors and prevent losses due to diffusion. A further complication is that the goods to be exchanged might often vary in their diffusion properties. Our assays, for instance, suggest that the diffusion and sharing of TasA is rather limited compared to EPS. We argue that such low diffusion rates must be compensated by an increased spatial mixing of the cooperation partners. Therefore, in opposition to “xenophobic” mechanisms employed by microbes to avoid strangers [48–50], “xenophilic” strategies might be crucial for genetic division of labor.

In conclusion, our study offers major insights into the evolution of division of labor. First, it shows that genetic specialization can be superior over phenotypic division of labor because it enables to break metabolic and regulatory constraints prevailing in organisms that remain totipotent. Second, sophisticated genetic division of labor can occur in simple organisms such as bacteria. Finally, genetic division of labor, based on the reciprocal exchange of public goods, could represent an evolutionary stable strategy, with strain frequency evolving towards an equilibrium that maximizes group productivity. Important to consider is whether de novo mutations may occur in the long term and disturb the observed equilibrium. For instance, a double mutant Δ*eps*Δ*tasA*, which is deficient in both matrix components could exploit the complementing partners and derail the genetic division of labor. Future studies will need to experimentally test whether the reported cases of genetic division of labor are evolutionary stable in the long run.

## Author Contributions

A.D. and Á.T.K. conceived the project, A.D., H.K., M.M., C.-Y.H., and C.E. performed experiments, R.H. and K.D. analyzed quantitatively the CLSM imaging data, C.-Y.H., N.S.-W., C.E. and S.B. analyzed the flow cytometry results, T.W. and R.K. performed modeling and analyzed the simulations. A.D., R.K., and Á.T.K. wrote and corrected the manuscript. All authors contributed critically to the drafts and gave final approval for publication.

## Acknowledgement

This work was funded by the Deutsche Forschungsgemeinschaft (DFG) to Á.T.K. (KO4741/2.1) within the Priority Program SPP1617. A.D. was supported by a fellowship from Alexander von Humboldt foundation. R.K. was funded by the Swiss National Science Foundation (grant no. PP00P3_165835) and the European Research Council (ERC-CoG no. 681295). This work was further supported by BBSRC grant code BB/P0001335 to N.S.-W., a scholarship from BeautyHsiao Biotech. Inc to C-Y.H., the European Research Council (ERC-StG no. 716734) and the Human Frontier Science Program (CDA00084/2015-C) to K.D., and a Start-up grant from the Technical University of Denmark to Á.T.K. Work in the laboratory of Á.T.K. is partly supported by the Danish National Research Foundation (DNRF137) for the Center for Microbial Secondary Metabolites. We acknowledge the help of Dr. Rosemary Clarke for assistance with flow cytometry performed at the University of Dundee.

## STAR*METHODS

### CONTACT FOR REAGENT AND RESOURCE SHARING

Further information and requests for resources and reagents should be directed to and will be fulfilled by the Lead Contact, Ákos T. Kovács.

### EXPERIMENTAL MODEL AND SUBJECT DETAILS

All bacterial strains used in this study derived from *Bacillus subtilis* NCBI 3610 *comI*^Q^12I^^ strain (Konkol et al., 2013). Strains were maintained in LB medium (Lysogeny broth (Lennox); Carl Roth, Germany), while MSgg medium was used for pellicle formation assay [1].

## METHOD DETAILS

### Strain construction

All strains that were used in this study or that were used solely as gDNA donors are listed in Table S1. To obtain TB601 and TB863, the NCBI 3610 *comI*^*Q12I*^ was transformed with gDNA isolated from DL1032 selecting for Tet-resistant colonies or Km-resistant colonies, respectively. TB524.1 and TB525.2 were obtained by transforming TB601 with gDNA isolated from TB500.1 and TB501.1, respectively. TB538.1 and TB539.1 were obtained by transforming TB602 with gDNA isolated from TB500.1 and TB501.1, respectively. To obtain TB864 and TB865, NCBI 3610 *comI*^*Q12I*^ was first transformed with gDNA from 168hymKate and then with gDNA isolated from NRS2242 and NRS3913, respectively. To obtain Anc Kate P_*eps*_-GFP, strain TB602 was first transformed with gDNA from 168hymKate and then with gDNA from NRS2242. To obtain Anc Kate P_*eps*_-GFP, strain TB 601 was first transformed with gDNA from 168hymKate and then with gDNA from NRS2394. In order to construct pTB848 and pTB849, the *eps* and *tapA* promoters were amplified using oTB172-oTB173 and oTB174-oTB175 primers pairs, respectively (see Table S2), the PCR products were digested with *Eco*RI and *Nhe*I, and cloned into the corresponding sites of vector pmKATErrnB. To obtain strains TB961 and TB962, first NCBI 3610 *comI*^*Q12I*^ was transformed with gDNA from NRS2242, and the obtained strain (TB373) was transformed with plasmids pTB848 and pTB849, respectively. TB960 was constructed by transforming NCBI 3610 *comI*^*Q12I*^ with gDNA from NRS3913 and the obtained strain (TB363) was subsequently transformed with pTB849 plasmid. To construct plasmid pTB498 harbouring a constitutively expressed mKATE2 gene, the P_hyperspank_-mKATE2 fragment was PCR amplified with primers oTH1 and oTH2 from plasmid phy-mKATE2 [2], digested with *Xba*I and *Eco*RI, ligated into plasmid pWK-Sp as described in [3]. Resulting plasmids were verified by sequencing and transformed into *B. subtilis* NCBI 3610 *comI*^*Q12I*^, resulting in TB539.

Plasmid pNW725 was used to construct strain NRS3913. This was generated through amplification of the *mKate2* coding region from plasmid pTMN387 using primers NSW1026 and NSW1027 (see Table S2) and ligation into plasmid pNW600 using *HinD*III and *Bam*HI. Plasmid pNW600 carries the P*tapA* promoter region (Murray et al., 2009), and therefore plasmid pNW725 has the *mKate2* coding region under the control of the *tapA* promoter region. Plasmid pNW725 was integrated into the chromosome of *B. subtilis* NCIB3610 at the *amyE* locus. Strain NRS5832 was generated by phage transduction of the P*epsA-gfp* reporter fusion from strain NRS2242 into NRS3913 as the recipient. Phage transduction was performed using SSP1 phage as previously described (Verhamme et al., 2007).

### Pellicle formation and productivity assays

To obtain pellicle biofilms, bacteria were routinely growth in static liquid MSgg medium at 30°C for 48 hours, using 1% inoculum from overnight cultures. Productivities where accessed by examining colony forming units (CFUs) in mature pellicles. Prior each CFU assays, pellicles were sonicated according to a protocol optimized in our laboratory that allows proper disruption of biofilms without affecting cell viability [4,5]. To access relative frequencies of Δ*eps* and Δ*tasA* strains, the cocultures were plated on selective antibiotics tetracycline (10μg/ml) and spectinomycin (100μg/ml), respectively.

### Fitness assays

Since the expression of *epsA-O* and *tapA-sipW-tasA* operons strongly depend on cultivation conditions and media composition [3,6–9], we performed the competition experiment for the fitness costs of EPS and TasA production under the same conditions that were later used for the assays that involved pellicles. Strains of interest were premixed at 1:1 ratios based on their OD_600_ values and the mixture was inoculated into MSgg medium at 1%. Cultures were grown under static conditions at 30°C. CFU assays (using selective antibiotics for the Δ*eps* and Δ*tasA* strains) were performed immediately after inoculation and after 16 hours of growth. The growth curves obtained at the initial stage of pellicle formation were performed under standard pellicle growth conditions in 96-well plates. The optical densities and GFP-fluorescence were monitored using an infinite F200PRO plate reader (TECAN Group Ltd, Männedorf, Switzerland).

### Spent media complementation assay

The supernatants were obtained from the WT, Δ*eps* and Δ*tasA* strains grown under static conditions in MSgg medium at 30°C for 48 hours. Cells were pelleted by centrifugation (5min, 8000 r.p.m.), the supernatants were sterilized using Millipore filters (0.2μm pore size), and mixed in 1:1 ratio with 2 times’ concentrated MSgg medium. Surface colonization of the Δ*eps* and Δ*tasA* in presence of conditioned media from the WT or complementary mutant strains were compared with the negative controls where the mutants grew in presence of their own conditioned media.

### Microscopy/confocal laser scanning microscopy (CLSM)

Bright field images of whole pellicles and colonies were obtained with an Axio Zoom V16 stereomicroscope (Carl Zeiss, Jena, Germany) equipped with a Zeiss CL 9000 LED light source and an AxioCam MRm monochrome camera (Carl Zeiss). For time-lapse experiment, cultures were grown in 24-well plates (1.5 cm diameter per well), incubated in INUL-MS2-F1 incubator (Tokai Hit, Shizuoka, Japan) at 30 °C and images were recorded every 15 min. The detailed description of the fluorescence time lapse microscope has been previously published [10]. The pellicles were also analyzed using a confocal laser scanning microscope (LSM 780 equipped with an argon laser, Carl Zeiss) and Plan-Apochromat/1.4 Oil DIC M27 63× objective. Fluorescent reporter excitation was performed with the argon laser at 488 nm and the emitted fluorescence was recorded at 484–536 nm and 567–654 nm for GFP and mKate, respectively. To generate pellicle images, Z-stack series with 1 μm steps were acquired. Zen 2012 Software (Carl Zeiss) was used for both stereomicroscopy and CLSM image visualization.

### Sample fixing and flow cytometry

Pellicles were harvested at 24, 48, and 72 h into sterile 2 ml screw cap tubes, followed by centrifugation at 17000′g for 10 min. GTA buffer (50 mM glucose, 10 mM EDTA pH 8.0, and 20 mM Tris-HCl pH 8.0) was added into 24-well plates to harvest the cells remained in wells and pooled with cell pellet from previous step. Pooled cell pellets were then pumped through 23G needles 6 times to disperse pellicles. Dispersed samples were pelleted down and fixed by incubation with 4% paraformaldehyde for 7 min at room temperature. Fixed samples were washed with GTA, and subjected to mild sonication prior flow cytometry. Flow cytometry (LSRFortessa^(tm)^, BD biosciences) were operated by FACS facility in School of Life Sciences, University of Dundee. For initial experiments comparing the expression of matrix genes in the WT and the biofilm mutants flow cytometry (BD Facscanto II, BD biosciences) was performed at Disease Systems Immunology Group, DTU Bioengineering.

### Root colonization assay/root biofilms productivity

Colonization of *Arabidopsis thaliana* roots was performed according to modified protocol from [7]. Arabidopsis ecotype Col-0 seeds were surface sterilized using 2% (v/v) sodium hypochlorite solution as follows: seeds were incubated in 2% (v/v) sodium hypochlorite with mixing on an orbital shaker for 20 min and then washed five times with sterile distilled water. The seeds were placed on pre-dried MS agar plates (Murashige and Skoog basal salts mixture; Sigma) (2.2 g l^-1^) in an arrangement approximately 20 seeds per plate at a minimum distance of 1 cm. Seeds were germinated and grown on agar plates containing MS medium. After 3 days of incubation at 4°C, plates were placed at an angle of 65° in a plant chamber (21°C, 16h light per day). After 6 days, homogenous seedlings ranging 0.8-1.2cm in length were selected for root colonization assay. Seedlings were transferred into 48-well plates containing 270μl of MSNg medium [7]per well. Next the medium was supplemented with 30μl of exponentially growing bacterial culture diluted to OD_650_ = 0.2. The sealed plates were incubated at rotary shaker at 28°C for 18h at 90 r.p.m. After the incubation, plants were washed 3 times with MSNg to remove non-attaching cells and then transferred to a glass slide for imaging using CLSM. To access root biofilm productivities, the roots were transferred into Eppendorf tubes, subjected to standard sonication protocol and the CFU assays were performed for obtained cell suspensions. To extract CFU/mm of root, the obtain CFU values were divided by total length of a corresponding root.

### Images of plant roots

For biofilm roots visualization, the GFP and mKate images were converted into 3D projections, contrast was enhanced using normalized function and green and red lookup tables were applied for GFP and mKate channels, respectively. Overlay images were obtained in ZEN software and further processed using ImageJ as follows: Brightness and contrast were adjusted, the root and biofilm area was manually selected and the background was lightened and smoothed using ‘adjust brightness’ and ‘smooth’ functions, respectively.

### Modelling

We performed individual-based simulations, using the platform developed by Dobay et al. (2014). Microbial simulations occur on a two-dimensional toroidal surface with connected edges (i.e. there are no boundaries). The surface of the torus is 10,000 μm^2^(100 × 100 μm). Bacteria are modeled as discs with an initial radius of 0.5 μm. Bacteria can consume resources, grow at a basic growth rate (μ = 1) and divide when reaching the threshold radius of 1 μm. In our simulations, we assumed that resources are not limited. Bacteria further produce beneficial public goods at a cost *c* per molecule and at constant rate of 1 molecule/s. Public goods diffuse randomly according to the diffusion coefficient *d* (μm^2^/s) and following a Gaussian random walk. Public goods can decay with a certain probability *p*, with *p* increasing exponentially with time following the exponential function *p* = 1 – *e* ^−*w*Δ*t*/∂^, where Δ*t* is the age of the molecule, ***w*** the stiffness of the decay and ∂ the durability of the molecule. A public good can generate a benefit *b* to the cell that takes it up, which occurs when the cell and the public good physically overlap on the landscape. Bacteria can randomly disperse, too, defined by the diffusion coefficient *D* (μm^2^/s). Because we aimed to model bacterial performance in biofilms, where cell dispersal is relatively low, we set *D* = 0.01 μm^2^/s. Important to note is that neither bacteria nor public goods are bound to a grid, but move on a continuous landscape (following an off-lattice model with double-precision numbers). This mimics natural bacterial behavior as close as possible. One practical complication of this approach is that cells overlap with each other following diffusion. To cope with this issue, we applied an overlap correction after each time step following the procedure described in [11].

Using this setup, we simulated the performance of a wildtype (WT) strain, producing two public goods representing EPS and TasA, and two strains (PG1 and PG2) producing only one of the two public goods. We arbitrarily considered PG1 = TasA producer and PG2 = EPS producer. The growth of the three strains is defined by the following recursive functions:

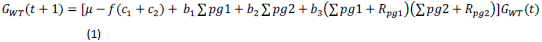

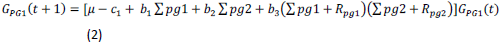

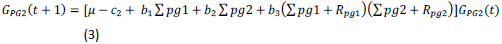

where *G* is the radius increase per time step *t*, μ is the basic growth rate, *c*_1_ and *c*_2_ are the costs of producing the respective public goods, and *f* is the metabolic constraint factor, whereby *f* > 1 if the simultaneous production of both public goods is costlier than producing either of the public goods alone. Furthermore, while *b*_1_ and *b*_2_ are the benefits accruing when a respective public good is taken up multiplied by the total number of public goods consumed (Σ*pg1* and Σ*pg2*) per time step, *b*_3_ is the synergistic benefit accruing for all the complementary public goods taken up within a certain period of time (*R*_*pg1*_ and *R*_*pg2*_, respectively). We arbitrarily chose five time steps for *R*_*pg1*_ and *R*_*pg2*_.

For all simulations, we seeded our in-silico landscape with eight cells placed in the center of the landscape to mimic the early phase of pellicle formation. Cells then started to produce public goods, grew and divided defined by their growth function. We let bacteria grow for 10,000 time steps in 50 independent replicates for each parameter combination. We examined three growth treatments, which included the WT strain in monoculture, the two complementary strains PG1 and PG2 in monocultures, and the two complementary strains PG1 and PG2 in mixed cultures. In the mixed cultures, we varied the starting frequency of the two strains from 1:7 (PG1 to PG2) to 7:1. For all simulations, we extracted the absolute productivity of the biofilm and the relative fitness of the competing strains within biofilms. To assess the role of public good diffusion on biofilm productivity and relative strain fitness, we varied public good diffusion from 3 to 7 μm^2^/s in steps of 0.5 μm^2^/s. To take into account that the public goods TasA and EPS might generate different benefits we varied the *b*_1_/*b*_2_ ratio from 1/9 to 1/1. Finally, we examined the effect of metabolic constraints on WT fitness by varying *f* from 1 to 1.3. All parameters together with the specific values used are given in the Supplementary Table S3.

## QUANTIFICATION AND STATISTICAL ANALYSIS

### Relative fitness

Relative fitness W_A_ for strain A in competition with strain B was calculated as follows:

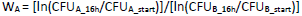

All replicates where one strain occurred to strongly dominate in the initial inoculum (exceeding initial 0.8 frequency) were removed from the dataset.

### Strain frequencies on plant roots

Ratios of the Δ*eps*^GFP^ and ΔtasA^mKate^ (and control with swapped fluorescent reporters) in root biofilms were estimated from the ratios of white pixel volumes measured on corresponding fluorescent images. Images were analyzed using ImageJ software. First, the root and biofilm area it was manually selected on the white-light image. For each channel, the stacks were converted into binary images and threshold was set up to > 0 value. Next, the root+biofilm selection was activated on the processed stacks and total pixel volumes for each channel were extracted using ‘stacks statistics’ function.

### Density correlation

The corresponding image stacks were dissected into cubes of 10 px side length. For each channel, the biovolume per cube was obtained. For all cubes containing biovolume in either of the two fluorescence channels (designated ch1 and ch2) the total biovolume in ch1 and ch2 within a sphere of a given radius (1 - 5 μm) was summed up, multiplied and normalized by the total volume of the sphere.

The resulting value ranges from 0 (no correlation, no biomass in one of the channels) over 0.25 (50% of biomass in ch1, 50% of biomass in ch2) to 1 (cube is completely filled in both channels = 100% overlap).

### Statistical analysis

For relative fitness assay, statistical differences from W=1 were identified using one-sample Student’s t-test. In case of productivity measurements statistical differences between two experimental groups were identified using two-tailed Student’s *t*-tests assuming equal variance. Variances in the two main types of datasets (relative fitness, productivity) were similar across different samples. No statistical methods were used to predetermine sample size and the experiments were not randomized. All relevant data are available from the authors.

## KEY RESOURCES TABLE

| Reagent or Resource | Source | Identifier |
| --- | --- | --- |
| **Chemicals** |  |  |
| Lysogeny broth (LB), Lennox | Carl Roth GmbH | X964.1 |
| Agar-Agar | Carl Roth GmbH | 5210.3 |
| Potassium Hydrogen Phosphate | Carl Roth GmbH | P749.1 |
| Potassium Dihydrogen Phosphate | Carl Roth GmbH | 3904.2 |
| L-Glutamic acid Monopotassium salt monohydrate | Alfa Aesar | 17232 |
| Magnesium chloride hexahydrate | Carl Roth GmbH | 2189.1 |
| Potassium chloride | Carl Roth GmbH | 6781.3 |
| Calcium chloride | Carl Roth GmbH | 5239.2 |
| Manganese(II) chloride | Carl Roth GmbH | T881.3 |
| Iron(III) chloride | Carl Roth GmbH | P742.1 |
| Zinc chloride | Carl Roth GmbH | T887.1 |
| Ammonium chloride | Carl Roth GmbH | K298.2 |
| Thiamin | Carl Roth GmbH | T911.1 |
| MOPS | Carl Roth GmbH | 6979.4 |
| Glycerol | Carl Roth GmbH | 7533.1 |
| Murashige and Skoog medium (MS) | Sigma Aldrich | M5519 |
| Sodium hypochlorite | Carl Roth GmbH | 9062.3 |
| Tetracycline hydrochloride | Carl Roth GmbH | 0237.1 |
| Spectinomycin dihydrochloride | Alfa Aesar | J61820 |
| Glucose | Fisher Scientific | G/0500/61 |
| EDTA | VWR | 20302.260 |
| Tris | VWR | 103157P |
| HCl | VWR | 20252.335 |
| **Experimental Models: Organisms/Strains** |  |  |
| *Bacillus subtilis* NCBI 3610 *comI*<sup>Q12I</sup> | [12] | DK1042 |
| *Bacillus subtilis* NCBI 3610 *comI*<sup>Q12I</sup> derivatives (listed in Table S2). | This study | N/A |
| *Arabidopsis thaliana* Col-0 | Greenhouse of Max Plank Institute for Chemical Ecology, Jena | N/A |
| **Recombinant DNA** |  |  |
| pmKATErrnB | [2] | Genbank Accession<br>number: KF245454 |
| pTB848 | This study | N/A |
| pTB849 | This study | N/A |
| pTB498 | This study | N/A |
| pNW725 | This study | N/A |
| **Sequence-Based Reagents** |  |  |
| Primers used in this study are listed<br>in Table S3. | This study | N/A |
| **Software and Algorithms** |  |  |
| ImageJ | [13] | https://imagej.nih.gov/ij/ |
| OriginPro 2015G | OriginLab, Northampton, MA | http://www.originlab.com/ |

## Supplementary Tables

**Table S1.**
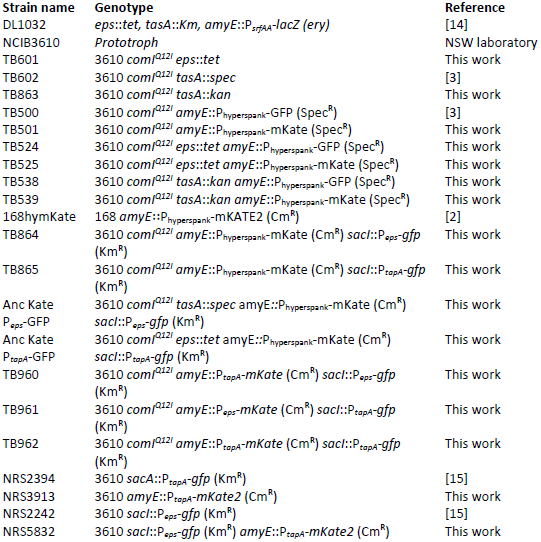
Bacterial strains used during experiments or as a source of genomic DNA

**Table S2.**
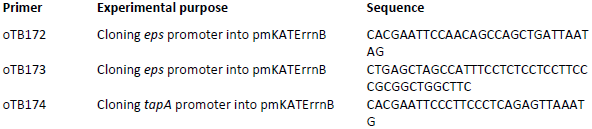
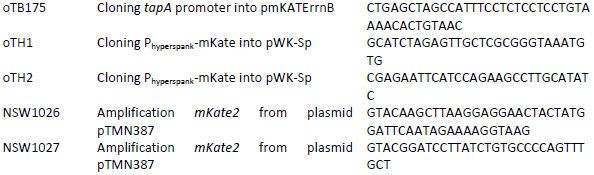
Primers used in this study

**Table S3.** Parameters and specific values used in modeling

| Parameter | Description | Value(s) |
| --- | --- | --- |
| $\mu$ | basic growth rate | 1 |
| $D$ | cell diffusion | 0.01 |
| $d$ | public good diffusion | 3 - 7 $\mu\text{m}^2/\text{s}$ |
| $\omega$ | stiffness of decay function | 0.1 |
| $\delta$ | public good durability | 500 s |
| $c_1$ | cost of public good 1 (TasA) | 0.0005 per molecule |
| $c_2$ | cost of public good 2 (EPS) | 0.0005 per molecule |
| $f$ | metabolic constraint factor | 1 - 1.3 |
| $b_1$ | benefit of public good 1 (TasA) | 0.0001 - 0.0009 |
| $b_2$ | benefit of public good 2 (EPS) | 0.0001 - 0.0009 |
| $b_3$ | synergistic benefit | 0.0005 |

