## Supplementary Materials for "Division of labor during biofilm matrix production"

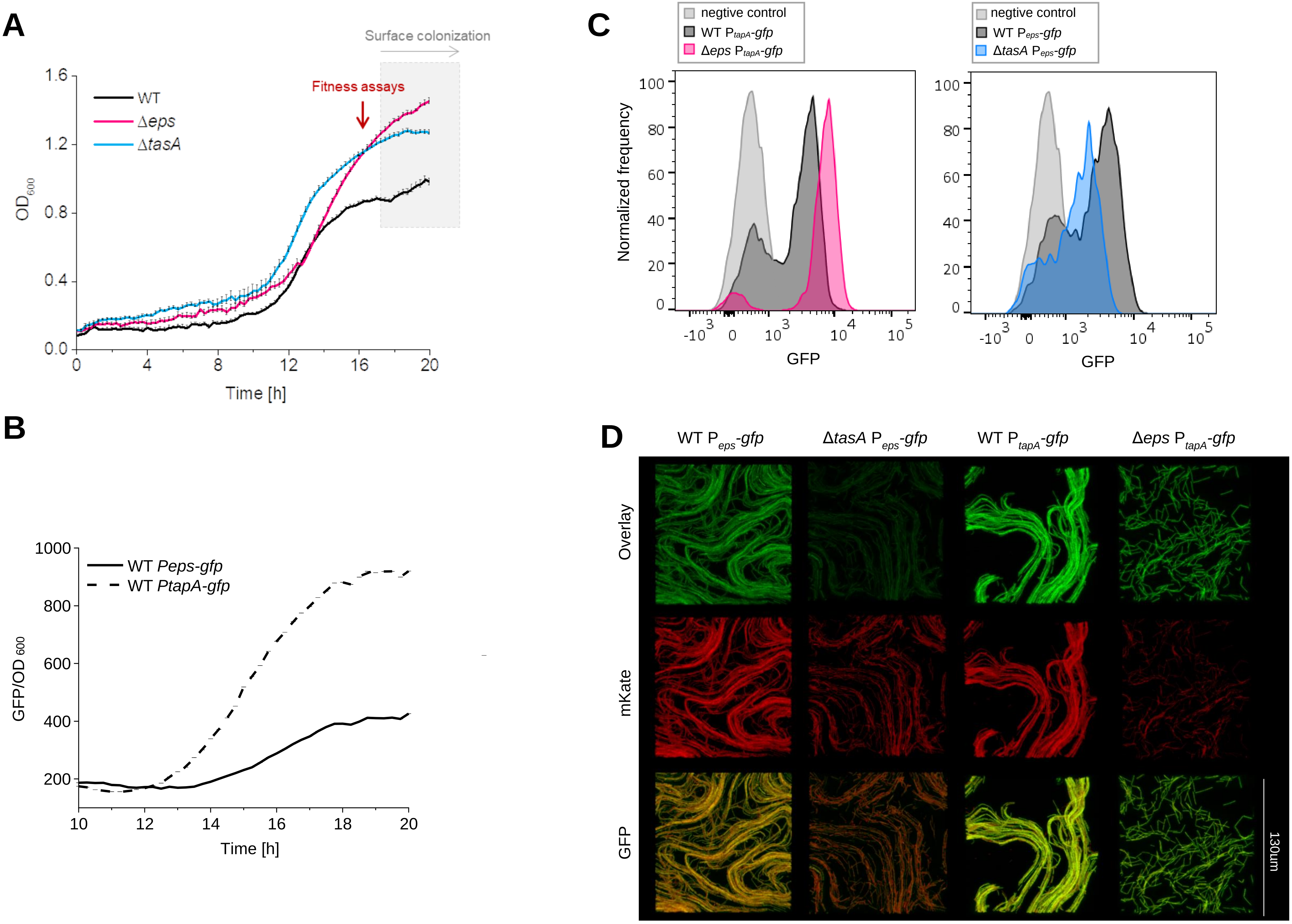
A
Surface colonization
Fitness assays
WT
Δeps
ΔtasA
OD600
Time [h]
B
WT Peps-gfp
WT PtapA-gfp
GFP/OD 600
Time [h]
C
negtive control
WT PtapA-gfp
Δeps PtapA-gfp
WT Peps-gfp
ΔtasA Peps-gfp
Normalized frequency
GFP
D
WT Peps-gfp
ΔtasA Peps-gfp
WT PtapA-gfp
Δeps PtapA-gfp
Overlay
mKate
GFP
130um

**Figure S1. Growth and matrix gene expression in the WT, Δ*eps* and Δ*tasA*. Related to Figure 1. (A)** Planktonic growth of the WT and matrix-deficient mutants Δ*eps* and Δ*tasA* were monitored for 20 h. Red arrow represents the time point selected as suitable for the fitness assays. Grey area corresponds to surface colonization phase where matrix components EPS and TasA become beneficial for the cells and when OD reads become less reliable due to cell clumping (see Movie S1). **(B)** Expression dynamics of *eps* and *tasA* were monitored in the WT using the $P_{eps}$-*gfp* and WT $P_{tasA}$-*gfp* reporter strains. It is important to note that the $P_{tapA}$-*gfp* reporter construct produced significantly stronger signal than the $P_{eps}$-*gfp*. Such differences can be due to actual differences in promoter strength or different efficiencies of promoter fusions. For panel A and B: data points represent an average (n=8) and error bars represent standard error. **(C)** Histograms of flow cytometric measurements for $P_{eps}$-*gfp* and PtapA-*gfp* in the WT and corresponding biofilm mutants. Y-axis shows normalized cell count and X-axis shows GFP fluorescence in arbitrary units. Negative control represents GFP expression in non-*gfp*-labelled WT strain. Representative images are shown for each strain (n=5). **(D)** Comparative analysis of CLSM images of Δ*eps* $P_{tapA}$-*gfp*, WT $P_{tapA}$-*gfp*, Δ*tasA* $P_{eps}$-*gfp* and WT $P_{eps}$-*gfp*, to evaluate the expression of remaining matrix genes (from $P_{tapA}$ and $P_{eps}$ promoters, respectively) by matrix-deficient strains. All strains used in this experiment constitutively expressed *mKate*. Width of each confocal image corresponds to 130 µm.

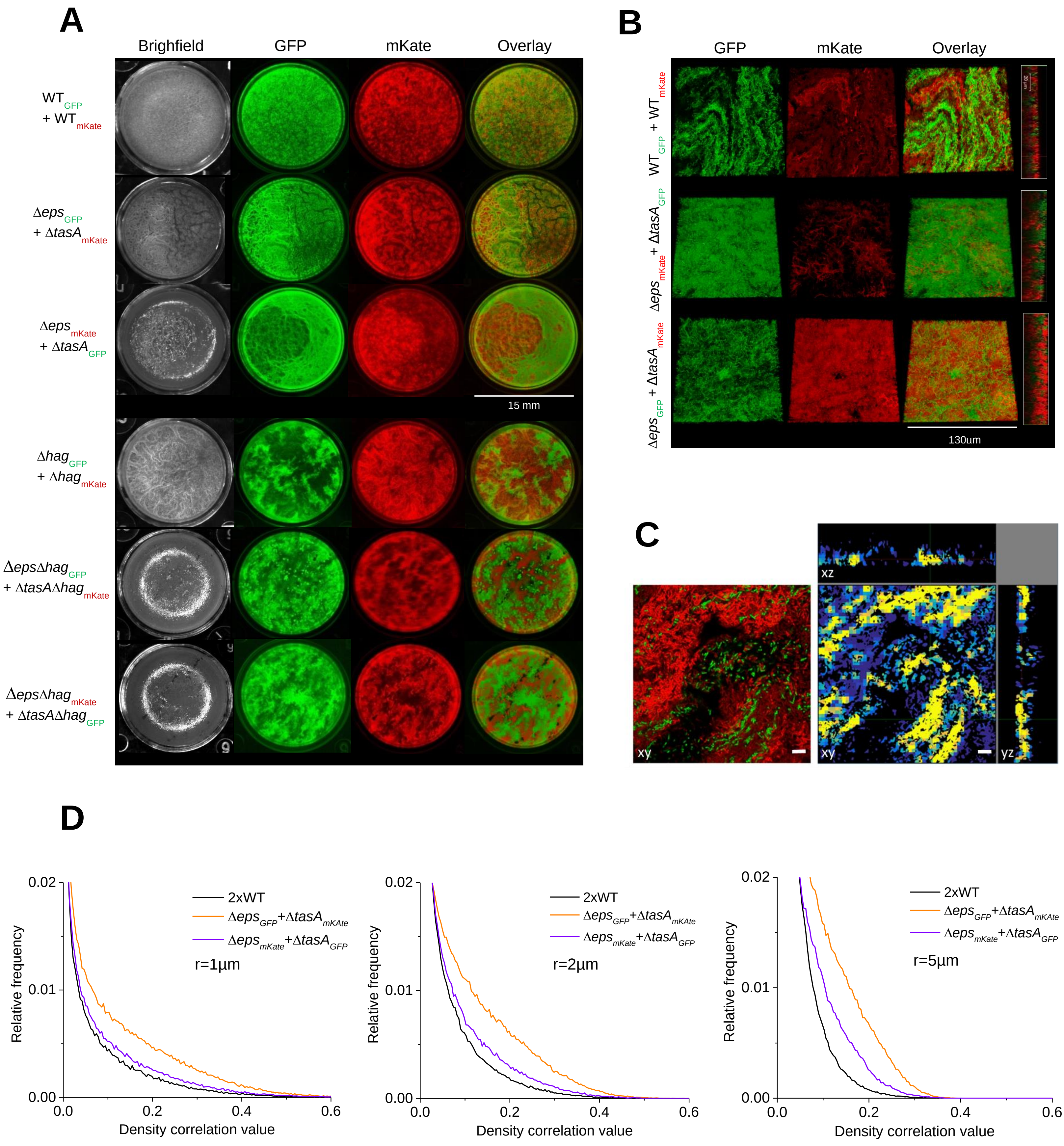

A
Brighfield
GFP
mKate
Overlay
WT GFP + WT mKate
Δeps GFP + ΔtasA mKate
Δeps mKate + ΔtasA GFP
15 mm
Δhag GFP + Δhag mKate
ΔepsΔhag GFP + ΔtasAΔhag mKate
ΔepsΔhag mKate + ΔtasAΔhag GFP
B
GFP
mKate
Overlay
WT GFP + WT mKate
Δeps mKate + ΔtasA GFP
Δeps GFP + ΔtasA mKate
130um
C
xz
xy
xy
yz
D
2xWT
Δeps GFP+ΔtasA mKAte
Δeps mKate+ΔtasA GFP
r=1µm
r=2µm
r=5µm
Relative frequency
Density correlation value
0.00
0.01
0.02
0.0
0.2
0.4
0.6

**Figure S2. Role of motility and mixing in complementation assay. Related to Figure 1. (A)** Three upper rows: matrix-proficient strains (WTs) and complementary mutant strains Δ*eps*+Δ*tasA* labelled with different constitutive fluorescent reporters were mixed in 50:50 ratios and allowed to form pellicles. Three bottom rows: the same experiment was performed in flagellum-deficient background (Δ*hag*). Assortment of strains in co-cultures was assessed after 48h (mature pellicles) using a fluorescence stereomicroscope. **(B)** Pellicles formed by WT strain mixtures with different fluorescent reporters (labeled 2xWT in the figure) and Δ*eps*+Δ*tasA* mixtures labelled with different constitutive fluorescent reporters were visualized using CLSM. **(C)** Visual representation of density correlation function applied in (D); Left: example pellicle image of $WT_{GFP}$+$WT_{mKate}$ obtained using CLSM; right: density correlation map for the same image with r = 2 µm. Blue: low density in either GFP or mKate channel. Yellow: high density in both channels. Scale bar represents 10µm. **(D)** Distribution of density correlation (2-class thresholding) for different ranges (see Methods) was compared for pellicles formed by matrix-proficient strains (WTs) and complementary mutants Δ*eps*+Δ*tasA*.

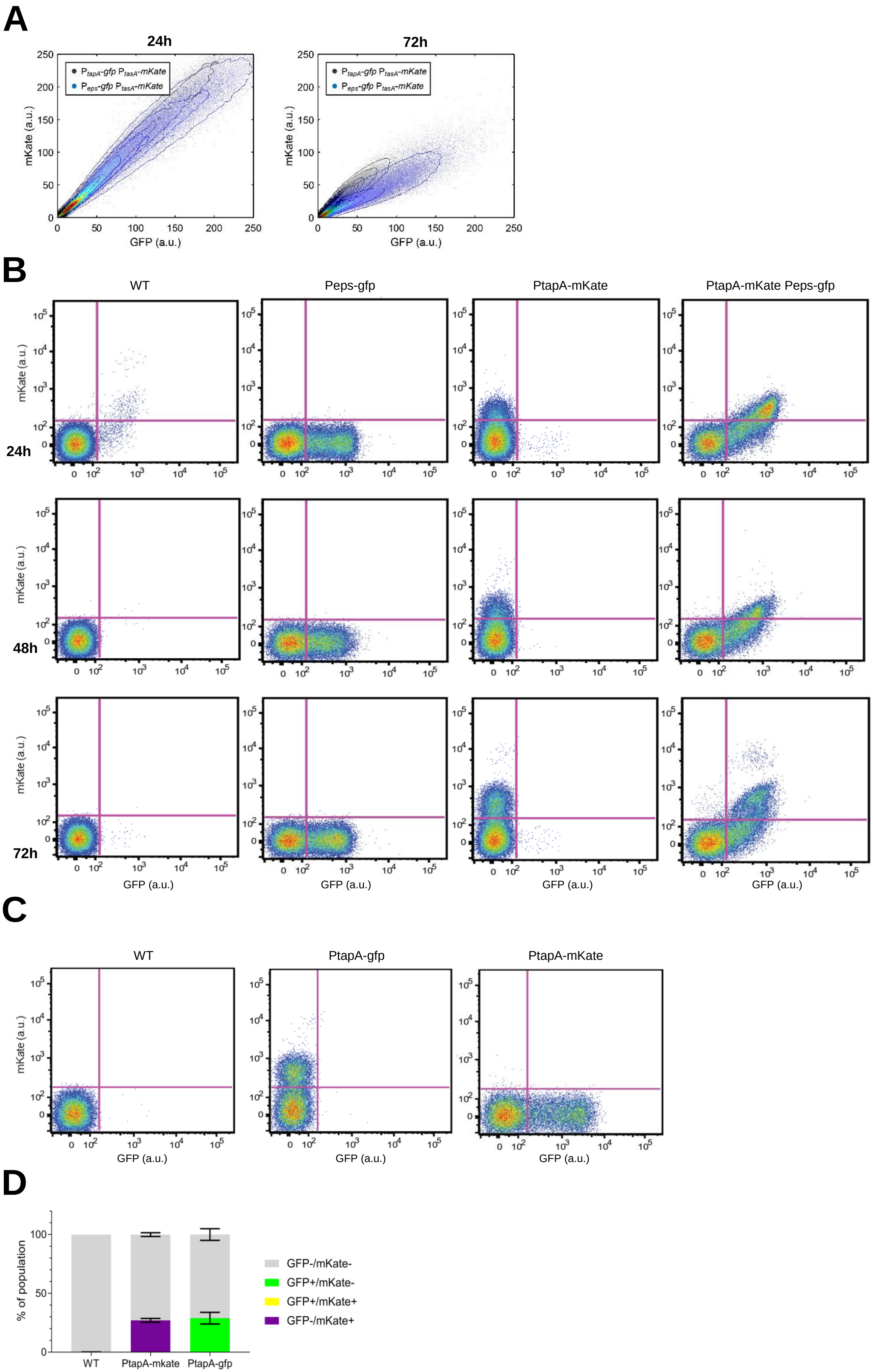

A
24h
72h
P_tapA-gfp P_tasA-mKate
P_eps-gfp P_tasA-mKate
mKate (a.u.)
GFP (a.u.)
B
WT
Peps-gfp
PtapA-mKate
PtapA-mKate Peps-gfp
24h
48h
72h
mKate (a.u.)
GFP (a.u.)
C
WT
PtapA-gfp
PtapA-mKate
mKate (a.u.)
GFP (a.u.)
D
% of population
GFP-/mKate-
GFP+/mKate-
GFP+/mKate+
GFP-/mKate+
WT
PtapA-mkate
PtapA-gfp

**Figure S3. Native phenotypic heterogeneity in the expression of matrix components accessed at different time points. Related to Figure 2. (A)** Volumes in GFP and mKate channels (obtained by manual thresholding) were merged, dissected into cubes and the average intensities in GFP and mKate for all cubes are plotted (see methods). The pellicles were sampled after 24h and 72h. The maximum density is normalized to 1 and the contour lines correspond to 0.05 decrease in density. **(B)** Pellicles formed by NCIB3610, NRS2242 ($P_{eps}$-*gfp*), NRS3913 ($P_{tasA}$-*mKate*) and NRS5832 ($P_{eps}$-*gfp* $P_{tasA}$-*mKate*) were sampled after 24h, 48h and 72h and analysed by flow cytometry. GFP (X-axis) and mKate (Y-axis) intensity were shown and threshold for fluorescent expression were set based on the cell distribution of NCIB3610. **(C)** Cell fluorescence distribution of NCIB3610, NRS3913 ($P_{tasA}$-*mKate*), and NRS2394 ($P_{tasA}$-*gfp*) obtained via flow cytometry. Pellicles were grown for 48 h. **(D)** Corresponding bar chart (mean±SD) representing the fractions of ON and OFF cells in the pellicles analyzed in (C) (n=2).

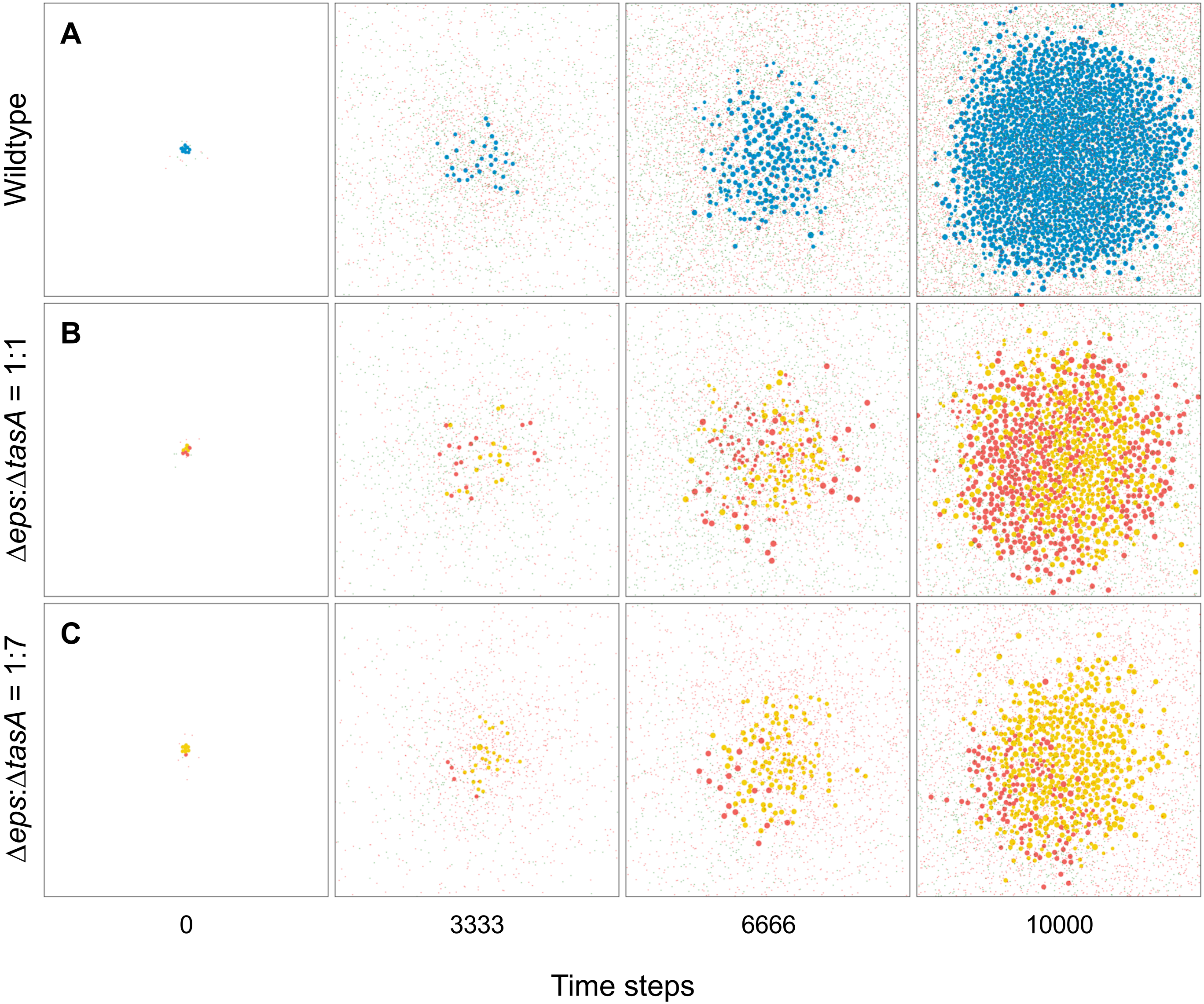
A
B
C
Wildtype
Δeps:ΔtasA = 1:1
Δeps:ΔtasA = 1:7
0
3333
6666
10000
Time steps

**Figure S4. Illustrative examples of simulated pellicle formation. Related to Figure 5. (A)** Wildtype bacteria (blue discs), simultaneously producing the two complementary public goods TasA and EPS (depicted by the tiny colored dots) grows to high cell densities and forms circular pellicles. **(B)** Co-culture of the two specialized mutants Δ*eps* (red discs, producing TasA) and Δ*tasA* (yellow discs, producing EPS) can complement each other by exchanging public goods. However, cell density is lower at the end of the simulation compared to the wildtype since these examples assume that there are no metabolic constraints of producing two public goods in the wildtype. **(C)** Inefficient complementation between the two mutants at a non-optimal strain ratio leads to low cell density in the pellicle. These examples further demonstrate that a high proportion of public goods is lost by diffusion, especially at low cell densities (i.e. early time points). Parameter settings were: public good diffusion $d = 5$, metabolic constraints $f = 1$, benefits $b_1 = b_2 = b_3 = 0.0005$.

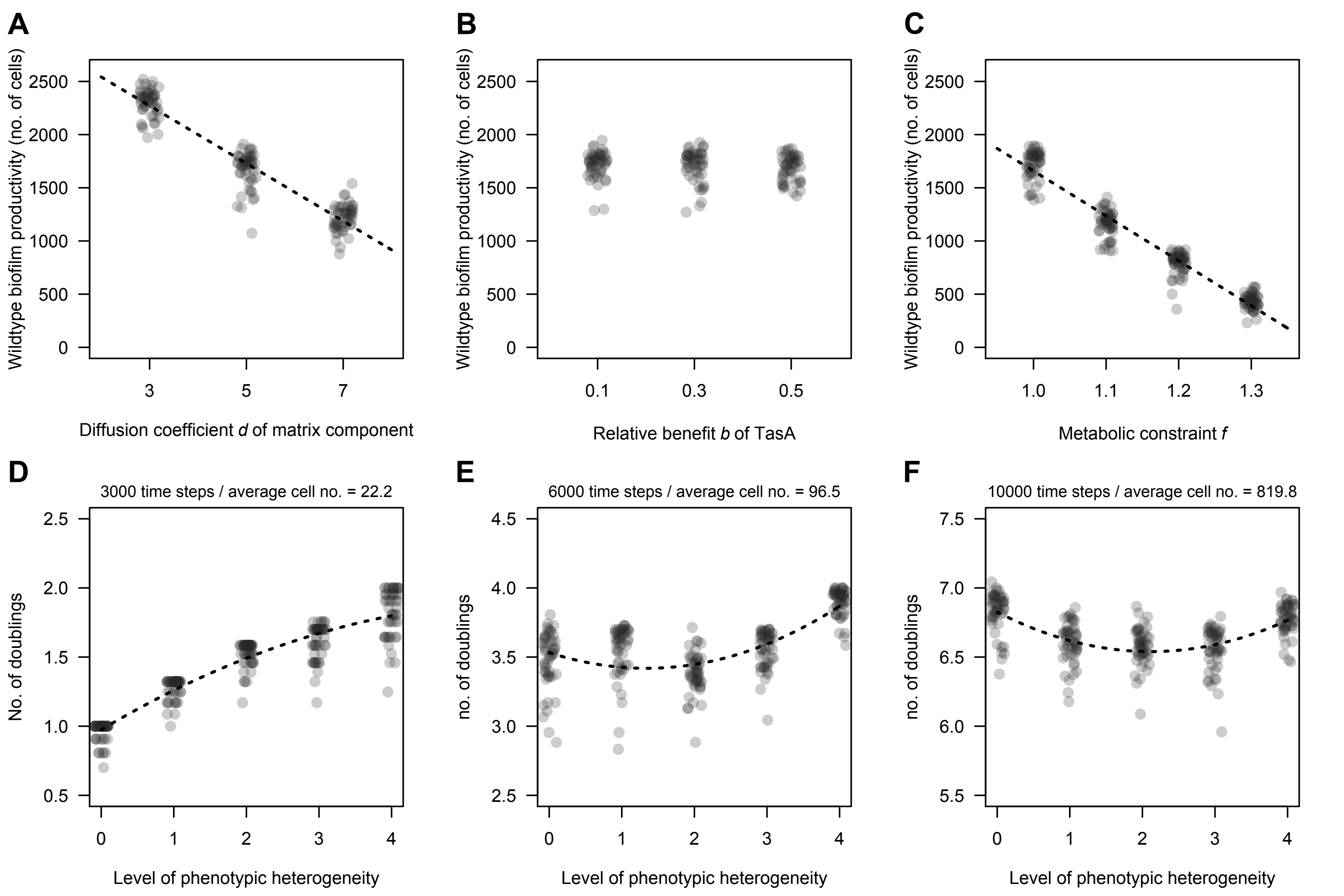
A
Wildtype biofilm productivity (no. of cells)
2500
2000
1500
1000
500
0
3
5
7
Diffusion coefficient d of matrix component
B
Relative benefit b of TasA
0.1
0.3
0.5
C
Metabolic constraint f
1.0
1.1
1.2
1.3
D
3000 time steps / average cell no. = 22.2
No. of doublings
2.5
2.0
1.5
1.0
0.5
0
1
2
3
4
Level of phenotypic heterogeneity
E
6000 time steps / average cell no. = 96.5
no. of doublings
4.5
4.0
3.5
3.0
2.5
F
10000 time steps / average cell no. = 819.8
7.5
7.0
6.5
6.0
5.5

**Figure S5. Variables affecting *in-silico* biofilm productivity of the wild type strain. Related to Figure 5. (A)** - **(C)** Simulated biofilm formation of the wild type strain, simultaneously producing TasA and EPS. Biofilms were initiated with eight cells, each producing the two matrix components at the same rate and growing according to their fitness function (see STAR methods). Panels show endpoint productivity of the biofilms after 10,000 time steps, under conditions, where the diffusion coefficient *d* of the matrix components **(A)**, the relative benefit *b* of the two matrix components **(B)**, or the strength of metabolic constraints *f* for simultaneously producing two matrix components (**C**) were varied. Panels **(D)** - **(F)** show the number of cell doublings for different time intervals and for different levels of phenotypic heterogeneity. All simulations started with 8 cells. The metabolic constraint value was set to $f = 1.16$. With this value, unspecialized and completely specialized groups perform equally well across the entire growth cycle. It thus allows to compare how intermediate levels of specialization compare relative to these two extremes. The implemented levels of phenotypic heterogeneity were: 0 = 8 unspecialized cells; 1 = 6 unspecialized cells + 1 TasA specialist + 1 EPS specialist; 2 = 4 unspecialized cells + 2 TasA specialists + 2 EPS specialists; 3 = 2 unspecialized cells + 3 TasA specialists + 3 EPS specialists; 4 = 4 TasA specialists + 4 EPS specialists. Dashed lines indicate trend lines of the best significant fit to the data. If not indicated otherwise, parameter settings were $d = 5$, $f = 1$, $b_1 = b_2 = b_3 = 0.0005$.
